# Analysis of Phylogeny of *Agrocybe* Genus Based on Nucleotide Database in NCBI

**DOI:** 10.1101/2022.07.27.501799

**Authors:** Long Wang, Qingping Zhou, Lingli Li

## Abstract

The study of the phylogeny of species is a basic discipline for the effective protection, development, and utilization of their germplasm resources. Since this century, the fungi of the *Agrocybe* genus have become the most successful class of medium and high-grade edible and medicinal fungi that have been artificially domesticated and grown on a large scale, and have high commercial value. However, there are few reports on its germplasm resources. To clarify the genetic and developmental relationship at the genus level of the *Agrocybe* in the world. In this paper, based on the NCBI Nucleotide database(National Center for Biotechnology Information), taking the single gene ITS sequence as the research object, the T92 (Tamura 3-parameters) parameter model was used to calculate the genetic distance, and the neighbor joining method was used to construct the genus-level phylogenetic tree of the *Agrocybe*. The results showed that the phylogenetic tree constructed by 192 ITS single gene sequences of 43 species of the *Agrocybe* (including 31 known species and 12 undetermined species) in the NCBI Nucleotide database was roughly divided into 6 branches and belonged to different groups, and 17 undetermined species of the *Agrocybe* sp. was preliminarily identified. This study has a certain reference value for the genetic relationship among different species of *Agrocybe* all over the world, as well as the selection of the parent bank and the protection of germplasm resources of each species of the genus.

## 0. Introduction

Edible Fungi are a kind of ideal food with rich nutrition, delicious taste, and strong fitness. They are also one of the three major foods for humans. They also have high medicinal value and are recognized as high-nutrient health food. The *Agrocybe* genus, which was first named by French Agrocybe FAYOD in 1889 [1], belongs to the Basidiomycota, Agaricomycetes, Agaricales, and Strophariaceae according to the Dictionary of Fungi 10th edition [2]. the type species of the *Agrocybe* genus is *Agrocybe praecox* (Fr.)Fayod[3]. Modern medicine indicates that the fungi of the *Agrocybe* genus are rich in B vitamins, mineral elements, and anticancer fungal polysaccharides, The inhibition rate of mouse sarcoma^180^ and Ehrlich ascites cancer reached 80%-90%[4]. At the same time, the *Agrocybe* genus has become the most successful class of high-grade edible and medicinal fungi in wild artificial wild domestication and large-scale planting, with high commodity value, and is favored for its unique flavor, rich nutrition, and special pharmacological efficacy [5]. Edible fungi germplasm resources are the foundation of scientific research and product development. However, in recent years, due to environmental pollution, deforestation, over-picking, and other factors, The species and quantity of the *Agrocybe* genus are decreasing year by year, and the selection of fine strains of the *Agrocybe* genus in artificial cultivation is less. Therefore, the preservation of strain culture is of great significance to the gene preservation of the fungal germplasm resources of the *Agrocybe* genus, the relative stability of the genetic performance of excellent strains, and its scientific research and production.

At present, most of the research on the fungi of the *Agrocybe* genus focuses on the description of macroscopic characteristics, cultivation techniques, extraction of polysaccharide components, genetic analysis, and determination, etc[6–16]. Few reports have been reported on the diversity of its germplasm resources. Dai Yucheng *et al.* reported seven species of the *Agrocybe* genus in China [17]. Jin Xin *et al.* reported three new records of the *Agrocybe* genus in China [18]. According to the latest molecular biology research report, the *Agrocybe* genus is generally a composite population, in which *Agrocybe salicacola* and *Agrocybe chaxingu* should belong to independent species and be classified as *Agrocybe cylindracea*[19,20]. Currently, most studies on the germplasm resources of the *Agrocybe* genus focus on the observation and determination of morphological or agronomic traits, and there is a lack of molecular phylogenetic studies at the genus level. To clarify the genetic development at the genus level of *Agrocybe* sp., this study is based on the NCBI nucleotide database(National Center for Biotechnology Information)[21], which analyzes the genetic diversity of the germplasm resources of the *Agrocybe* genus throughout the world, providing a reference for in-depth research in the future.

## 1. Materials and Methods

### 1.1 Data source

Specific information on the *Agrocybe* genus was obtained using the NCBI-Taxonomy database. A total of 43 species information was retrieved, including 31 known species information and 12 pending species information. 192 ITS (Internal Transcribed Spacer, (ITS1-5.8S rRNA-ITS2) were retrieved and collected from the NCBI-Nucleotide database for analysis.

### 1.2 Data analysis

Perform homologous sequence alignment with Alignment-Clustal W of the collected *Agrocybe* genus ITS sequences by MEGA-X software, and the genetic distance was calculated using the Maximum Likelihood Method [22]. The Neighbor-Joining Method constructed the phylogenetic tree of each species in *Agrocybe* genus. The confidence level of the system branches was evaluated using a self-guided test, and the bootstrap support rate of each branch was set for 1000 repeated tests [23].

## 2. Results and analysis

### 2.1 Selection of statistical models

To improve the accuracy of phylogenetic tree construction, the maximum likelihood method was used in the MEGA-X software MODELS program to find the most appropriate genetic distance statistical model in Find the Best DNA / Protein Models, and the default parameters were selected during the analysis. Tab. 1 showed the maximum likelihood fit (including individual and combined models) of the 24 different nucleotide substitution models after systematic parameter statistics. In the composite model, The T92 +G combination model (BIC value =40706.37467) could be considered the best description; the AICc value of the T92 + G combined model was 36962.01904, The values were also relatively low, Slightly higher than the two combined models: GTR + G (AICc value =36944.80364) and GTR+ R + I (AICc value =36946.66629), This indicated that the combination model T92 + G had the highest overall fit degree, is the optimal solution out of the 24 substitution models. In the single model, the T92 model with BIC value =42122.97704 and AICc value =38388.3660, the lowest score and highest fit in the six single models. Because the MEGA-X version does not provide the phylogenetic tree construction of the combined model, the T92 model with the lowest BIC value was finally selected to build the phylogenetic tree at the genus level of the *Agrocybe* genus.

**Tab.1.** The maximum likelihood fit of 24 different nucleotide substitution models

| Model | parameter | Bayesian Information Criterion (BIC) | Akaike Information Criterion, corrected (AICc) | Maximum Likelihood value(lnL) | Gamma distribution |
| --- | --- | --- | --- | --- | --- |
| T92+G | 384 | 40706.37467 | 36962.01904 | -18095.84777 | 0.861605897 |
| HKY+G | 386 | 40729.80454 | 36965.95923 | -18095.80572 | 0.86051765 |
| T92+G+I | 385 | 40730.10236 | 36976.00188 | -18101.83313 | 0.858041331 |
| K2+G | 383 | 40735.96337 | 37001.35262 | -18116.5206 | 0.866323412 |
| GTR+G | 390 | 40747.62794 | 36944.80364 | -18081.20347 | 0.871034489 |
| TN93+G | 387 | 40748.08624 | 36974.49613 | -18099.06809 | 0.854771303 |
| HKY+G+I | 387 | 40753.86646 | 36980.27635 | -18101.9582 | 0.853004327 |
| GTR+G+I | 391 | 40759.23526 | 36946.66629 | -18081.12864 | 0.870953968 |
| K2+G+I | 384 | 40760.4191 | 37016.06347 | -18122.86998 | 0.862946708 |
| TN93+G+I | 388 | 40773.34096 | 36990.00609 | -18105.81696 | 0.84868148 |
| JC+G | 382 | 41011.34112 | 37286.47530 | -18260.08797 | 0.880813161 |
| JC+G+I | 383 | 41023.14393 | 37288.53319 | -18260.11089 | 0.880844977 |
| T92+I | 384 | 41788.70345 | 38044.34782 | -18637.01216 | n/a |
| K2+I | 383 | 41807.83225 | 38073.22151 | -18652.45505 | n/a |
| HKY+I | 386 | 41813.60269 | 38049.75738 | -18637.7048 | n/a |
| TN93+I | 387 | 41836.52594 | 38062.93584 | -18643.28794 | n/a |
| GTR+I | 390 | 41842.90036 | 38040.07607 | -18628.83968 | n/a |
| JC+I | 382 | 42089.27711 | 38364.41129 | -18799.05597 | n/a |
| T92 | 383 | 42122.97704 | 38388.36630 | -18810.02744 | n/a |
| K2 | 382 | 42147.39128 | 38422.52546 | -18828.11305 | n/a |
| HKY | 385 | 42148.81287 | 38394.71238 | -18811.18838 | n/a |
| TN93 | 386 | 42160.49446 | 38396.64914 | -18811.15068 | n/a |
| GTR | 389 | 42182.82942 | 38389.74982 | -18804.6827 | n/a |
| JC | 381 | 42429.16339 | 38714.04252 | -18974.87759 | n/a |

### 2.2 Construction of the phylo-genetic tree

Based on the T92 model and the neighbor-joining method, the ITS sequence phylogenetic tree of the *Agrocybe* genus was constructed (Fig.1). The clustering results showed that the entire phylogenetic tree was mainly divided into six major branches, covering 43 species of the *Agrocybe* genus in the NCBI-Taxonomy database (including 31 known species and 12 undetermined species), and each branch point had a high support rate (⩾ 0.572). The analysis showed that Agrocybe cylindracea (Agrocybe aegerita) and Agrocybe salicacacola were more closely related to each other, and they were clustered into one class with a topological value of 0.989, which could be classified as the sect. Aporus, the clustering results are consistent with the results of Zhang Jinxia et al[19]. Secondly, *Agrocybe erebia* with different accession numbers was clustered into a single group, with a clear phylo-genetic relationship, and together with *Agrocybe brunneola,* they were grouped into the Sect. Velatae. Thirdly, *Agrocybe pediades,* and their ring-shaped variety *Agrocybe pediades,* and *Agrocybe pediades* are grouped as a whole, and they are collectively classified as Sect. Pediades. In addition, *Agrocybe dura, Agrocybe paludosa,* and *Agrocybe praecox* are classified as Sect. Agrocybe. It was worth noting that the accession number MN007012.1 *Agrocybe parasitica* was relatively rare and was clustered with *Agrocybe cylindracea (Agrocybe aegerita)* and *Agrocybe chaxingu,* and its topological value = 0.501, which is worthy of specific genetic traits of *Agrocybe parasitica* of MN007012.1 to carry out in-depth research.

**Fig. 1.**
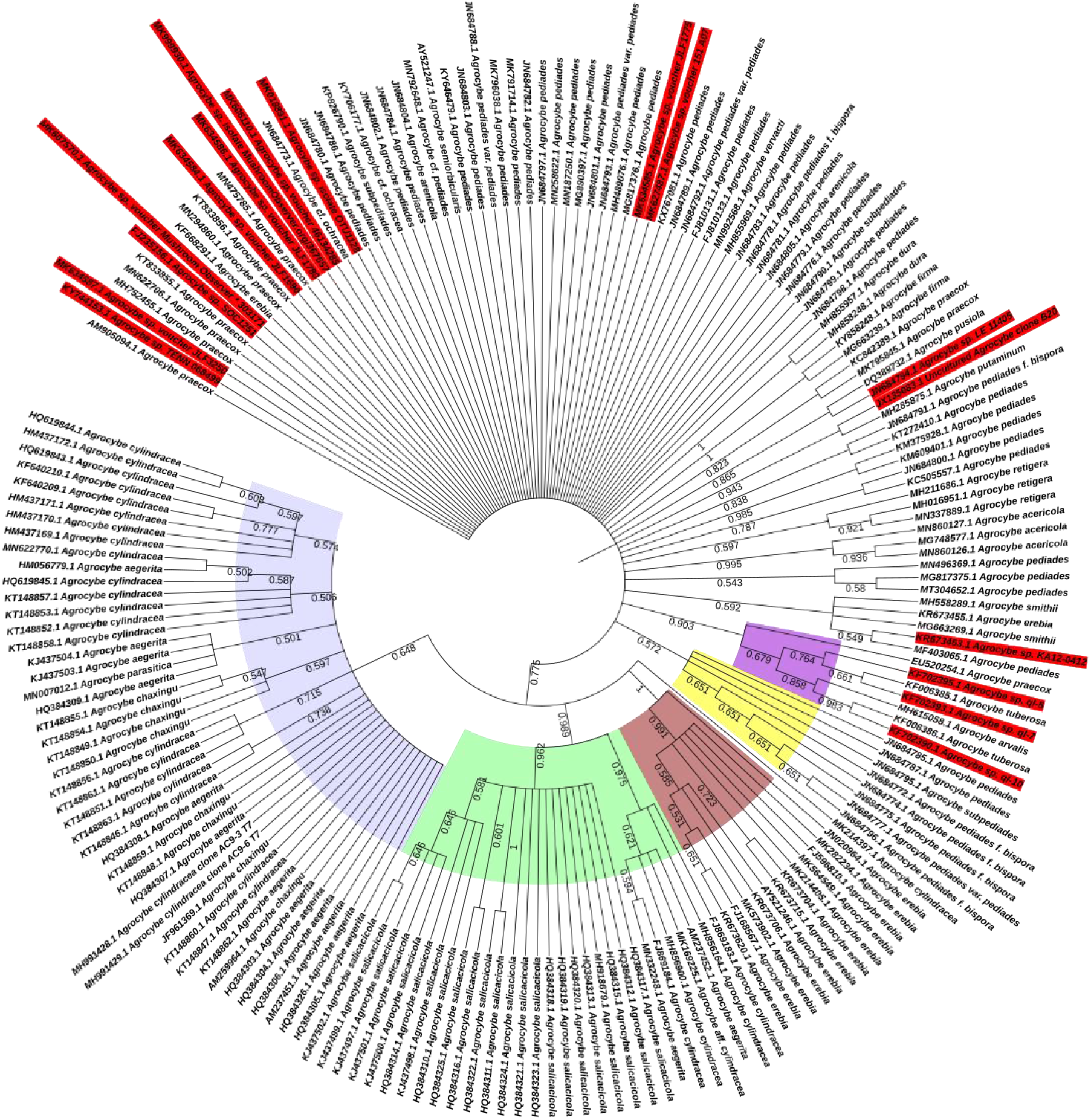
The phylogenetic tree of the *Agrocybe* genus based on NCBI-ITS sequence (Bootstrap, Repeat=1000)

Among the 192 ITS single gene sequence information recovered, 17 unknown species information were to be determined (marked on the red background). Among them, the accession number KY744153.1 (*Agrocybe* sp. TENN 068499), MK634587.1 (*Agrocybe* sp. voucher JLF3250), FJ235156.1 (*Agrocybe* sp. SOC1251), MK607570.1 (*Agrocybe* sp. voucher Mushroom Observer* 303171), MK634584.1 (*Agrocybe* sp. voucher JLF1690), MK634586.1 (*Agrocybe* sp. voucher JLF1780), MK999930.1 (*Agrocybe* sp. isolate Mushroom Observer.org/367 657) and MK606110.1 (*Agrocybe* sp. voucher 46134285) 8 undetermined species were clustered with the type-species *Agrocybe praecox*, and there is no branch point because they belong to parallel species. Accession numbers MK018891.1 (*Agrocybe* sp. isolate OTU1128), MK634585.1 (*Agrocybe* sp. voucher JLF1775), MK627487.1 (*Agrocybe* sp. voucher 151 A07), and KR673463.1 (*Agrocybe* sp. KA12-0412) 4 pending species were clustered with *Agrocybe pediades*, and also belonged to parallel species. Accession numbers KF702395.1 (*Agrocybe* sp. ql-5), KF702393.1 (*Agrocybe* sp. ql-7), and KF702390.1 (*Agrocybe* sp. ql-10) 3 undetermined species were clustered together with *Agrocybe tuberosa,* Topology value ⩾ 0.661. The accession number JX135083.1 (Uncultured Agrocybe clone B20) was clustered with *Agrocybe putaminum*, and the topological value = 0.9343, which has a very high similarity. The accession number JN684794.1 (Agrocybe sp. LE 11405) was clustered with *Agrocybe pusiola,* the topology value=0.865, and the similarity was also very high.

## 3. Conclusions

In this study, the ITS single gene sequences of each species of the *Agrocybe* genus were analyzed in the Nucleotide database of the National Center for Biological Information in the United States, and the genetic distance was calculated using the T92 model(Tamura 3-parameters). The neighbor-joining method constructed a phylogenetic tree of the *Agrocybe* genus. The results showed that the phylogenetic tree constructed by 192 ITS single gene sequences collected from the *Agrocybe* genus was roughly divided into 6 branches and belonged to different groups. Specific information is as follows: 8 pending species with accession numbers KY744153.1, MK634587.1, FJ235156.1, MK607570.1, MK634584.1, MK634586.1, MK999930.1, and MK606110.1 were classified in parallel with the type species *Agrocybe praecox.* The four pending species with accession numbers MK018891.1, MK634585.1, MK627487.1, and KR673463.1 also belonged to the parallel classification with A. pediades. The 3 undetermined species with accession numbers KF702395.1, KF702393.1 and KF702390.1 had a topological value of 0.661 with *Agrocybe tuberosa*, which was highly reliable. The topological value of accession number JX135083.1 and *Agrocybe putaminum* reached 0.943, and the similarity was extremely high, which could be considered the same species. The similarity of the JN684794.1 log-in numbers and *Agrocybe pusiola* similarity is also extremely high, with a topological value of 0.865, which can also be considered the same species. The accession number JN684794.1 was also very similar to *Agrocybe pusiola,* with a topological value of 0.865, which could also be regarded as the same species. In addition, special attention was paid to the accession number MN007012.1 *Agrocybe parasitica,* which had not yet been found in China. It was clustered with *Agrocybe cylindracea (Agrocybe aegerita)* and *Agrocybe chaxingu.* Topologically value reached 0.501, which belonged to a category with high reliability, and it was worthy of scholars to continue to carry out in-depth research on its specific genetic traits.

In 1994, Lu Chengying & Li Jianzong isolated a fungus of the Agrocybe genus in Zhangjiajie National Forest Park in Hunan Province, China, which was identified as *Agrocybe farinacea* Hongo by morphological characteristics, which was the newly recorded species in China at that time, specimen No. MHJSU940141, stored in Jishou University Fungal Herbarium (MHJSU)[24]. In 2012, Jin Xin and Tuliguer conducted a detailed systematic study on the taxonomy of domestic Agrocybe species, and counted three new recorded species in China, namely *Agrocybe elatella*, *Agrocybe brunneola,* and *Agrocybe pediades* var. cinctula Nauta, specimens of them were preserved in Jilin Agricultural University Herbarium (HMJAU)[25]. However, interms of the research by Chinese scholars on *Agrocybe farinacea*, *Agrocybe elatella,* and *Agrocybe brunneola*, the author did not find any information records for these three species in the NCBI-Taxonomy retrieval system, and this work should be improved in future research to enrich the NCBI strain resource bank.

For a long time, the taxonomy of the *Agrocybe* genus has been chaotic, and there is no unified classification standard. The classification research of this genus at home and abroad mainly includes traditional morphology, molecular systematics, and interspecies isolation-mating test. Modern molecular phylogenetic classification has become a powerful method for the study of fungal species classification, and a variety of molecular marker techniques have emerged[26,27]. Existing molecular phylogenetic studies of the *Agrocybe* genus mainly focus on phylogenetic studies of a single gene or two gene segments but cannot fully reflect its phylogenetic relationship. The construction of the phylogenetic tree of fungal multigene fragments has been adopted by the majority of scholars, which is more conducive to the correct definition of difficult species or similar species. However, the polygenic molecular phylogeny of the *Agrocybe* genus has not been reported. Therefore, the polygenic molecular phylogeny is one of the research hotspots in the molecular phylogeny of the genus Agrocybe. Due to environmental changes caused by geographical factors, the *Agrocybe* genus is unstable in morphology and physiological and biochemical characteristics and is prone to changes. At this time, mating experiments should be used to verify whether they produce reproductive isolation.

The phylogenetic study of species is the basic subject of biodiversity. The richer the genetic variation of a species, the stronger its survival adaptability and the greater its evolutionary potential. During its long-term natural evolution and artificial domestication, macrofungi have formed different ecological species to adapt to the ecological environment of different regions, and have relatively distinct genetic differentiation. This study has a certain reference value for the genetic relationship among different species of the *Agrocybe* genus, as well as the selection of the parent bank of each species and the protection of germplasm resources.

In recent years, with the rapid development of molecular sequencing technology, the Genealogical Concordance Phylogenetic Species Recognition (GCPSR) species identification system has become the main and hot spot in the field of biodiversity research, and there have been many research reports[28,29]. Through the rapid identification of more gene families, strains morphology, and reproductive mode, this technology can help humans understand the relatedness between species more effectively.

## Authors’ Contributions

Long Wang conceived and designed the experiments and translated the manuscript. Qingping ZHOU and Lingli LI performed the investigations and data analysis and wrote the original manuscript. All authors reviewed the manuscript.

## Acknowledgments

This research was supported by The Higher Education Institution Scientific Research Project in Hebei Province(ZD2020403); Doctoral Research Foundation of Tangshan Normal University (2020A13); Tangshan Agricultural Science and Technology Project(19150203E).

